# Floral presence and flower identity alter cereal aphid endosymbiont communities on adjacent crops

**DOI:** 10.1101/2023.01.11.523439

**Authors:** Sharon E. Zytynska, Sarah Sturm, Cathy Hawes, Wolfgang W Weisser, Alison Karley

## Abstract

1. Floral plantings adjacent to crops fields can recruit populations of natural enemies by providing flower nectar and non-crop prey to increase natural pest regulation. Observed variation in success rates might be due to changes in the unseen community of protective endosymbionts hosted by many herbivorous insects, which can confer resistance to various specialist natural enemies, e.g. parasitoid wasps. Reduced insect control may occur if highly protective symbiont combinations increase in frequency via selection effects, and this is expected to be stronger in lower diversity systems.
2. We used a large-scale field trial to analyse the bacterial endosymbiont communities hosted by cereal aphids (*Sitobion avenae*) collected along transects into strip plots of barley plants managed by either conventional or integrated (including floral field margins and reduced inputs) methods. In addition, we conducted an outdoor pot experiment to analyse endosymbionts in *S. avenae* aphids collected on barley plants that were either grown alone or alongside one of three flowering plants, across three time points.
3. In the field, aphids hosted up to four symbionts. The abundance of aphids and parasitoid wasps was reduced towards the middle of all fields while aphid symbiont species richness and diversity decreased into the field in conventional, but not integrated, field-strips. The proportion of aphids hosting different symbiont combinations varied across cropping systems, with distances into the fields, and were correlated with parasitoid wasp abundances.
4. In the pot experiment, aphids hosted up to six symbionts. Flower presence increased natural enemy abundance and diversity, and decreased aphid abundance. The proportion of aphids hosting different symbiont combinations varied across the flower treatment and time, and were correlated with varying abundances of the different specialist parasitoid wasp species recruited by different flowers.
5. *Synthesis and applications*. Floral plantings and flower identity can have community-wide impacts on the combinations of bacterial endosymbionts hosted by herbivorous insects. Our work highlights the potential of within-season selection for symbiont-mediated pest resistance to natural enemies with biological control impacts. This could be mitigated through increased recruitment of diverse natural enemies by incorporating functional diversity of floral resources into the environment.

## Introduction

Floral plantings are a common measure to recruit and establish natural enemies for natural pest control in adjacent crops under the framework of conservation biological control. However, inconsistent outcomes of floral plantings have hindered more widespread use (Albrecht *et al*. 2020). While we know a substantial amount about the effects of plant identity and diversity on insect herbivores and natural enemies of herbivores, we still lack understanding of how these interactions mediate, or are mediated by, other trophic groups such as insect-associated microbes, e.g. symbiotic bacteria (Hrcek, McLean & Godfray 2016). This has been particularly studied in plant-sucking insects such as aphids, which are major pests of crop worldwide (Zytynska, Tighiouart & Frago 2021). Since many aphid bacterial endosymbionts can also confer resistance to specialist natural enemies it follows that floral plantings that recruit a low diversity of natural enemies may also select for the optimal protective combination of microbes in the aphid host, with reduced biocontrol effectiveness (Zytynska & Meyer 2019).

The diversity of the aphid microbiome is surprisingly low (Sugio *et al*. 2015), with one obligate (primary), and nine common facultative (secondary) bacterial symbionts that have been identified in many different aphid species across the world (Zytynska & Weisser 2016). Despite the low diversity, these bacterial symbionts can have strong effects on aphid survival through providing essential nutrients and conferring resistance to natural enemies (parasitoid wasps and entomopathogenic fungi) or resistance to heat-stress, among other potential benefits (Oliver, Smith & Russell 2014; Guo *et al*. 2017). However, aphids experience fecundity and longevity costs of hosting symbionts, with varying costs associated with multiple hosting of different symbiont species (Zytynska, Tighiouart & Frago 2021). Furthermore, effects are highly aphid and symbiont species (and strain/genotype), specific leading to highly variable outcomes. A recent meta-analysis highlighted this variation and the bias towards information on a few common model aphid species (*Acyrthosiphon pisum, Sitobion avenae and Aphis fabae*) (Zytynska, Tighiouart & Frago 2021). Briefly, across multiple aphid species, endosymbionts *Hamiltonella defensa and Regiella insecticola* show strong protective effects against parasitism by parasitoid wasps. Other protective effects were supported by too few studies to be assessed in a meta-analysis, but include resistance to entomopathogenic fungi by *R. insecticola, Rickettsia, Ricketsiella viridis, Spiroplasma* and *Fukatsuia symbiotica*; parasitoid resistance by *Serratia symbiotica, F. symbiotica*; and heat-shock resistance by *S. symbiotica, H. defensa, Rickettsia*, and *F. symbiotica* (reviewed in Guo *et al*. (2017). There is also some evidence that infection with *H. defensa* can alter aphid anti-predator behaviour against generalist predators such as ladybirds (Polin, Simon & Outreman 2014; Humphreys, Ruxton & Karley 2022). Generally, field-collected aphids have been found to host 1-4 symbionts per individual (Zytynska & Weisser 2016), leading to high chance of aphids co-hosting multiple symbionts with variable costs and benefits. Furthermore, ineffective vertical transmission from mother to offspring (Rock *et al*. 2018) and variable selection pressures over time (e.g. changing natural enemy abundances, Smith *et al*. (2015)), as well as incompatible symbiont combinations in the field (Oliver, Moran & Hunter 2006) result in a polymorphism of infection within populations even in the same host-plant patch (Russell *et al*. 2013; Smith *et al*. 2015; Zytynska *et al*. 2016). Dynamic aphid-symbiont frequencies in a population will feedback to the interacting natural enemy communities with community-wide consequences and potential impacts on biological pest control efforts (Zytynska & Meyer 2019).

The impact of protective bacterial symbionts on plant-insect food webs, the regulation of aphid populations, and predator-prey dynamics is only beginning to be considered (McLean *et al*. 2016; Simon, Biere & Sugio 2017; Preedy *et al*. 2020; Carpenter *et al*. 2021; Leclair *et al*. 2021). It was previously shown that as plant species richness increased along a gradient (1-4 species, created as mixed plant communities from a pool of 20), so did the number and diversity of aphid symbionts identified in three aphid species (each exclusively feeding on a different host plant) in a long-term grassland experiment (Zytynska *et al*. 2016). One hypothesis is that that this is driven by a diversification in the number of selection pressures on the aphids (reviewed in Zytynska & Meyer 2019), through increased natural enemy diversity; a previous study in the long-term grassland experiment showed that as plant species richness increased so did parasitoid wasp diversity (Petermann *et al*. 2010). Therefore, when natural enemy diversity is low (e.g. in crop monocultures) there may be one optimal combination of symbionts that protects against a single main selection pressure (i.e. a dominant natural enemy). If all surviving aphids host the highly protective symbionts, then the diversity of symbionts in the population will be low (all aphids host similar symbiont communities). In a diverse plant community, with higher natural enemy diversity (including various parasitoid wasp species, entomopathogenic fungi and generalist predators), then there may be no single optimal symbiont combination. In this case, variable costs and benefits of co-hosting multiple symbionts could lead to aphids hosting various combinations of the available symbionts (higher diversity) (Hafer-Hahmann & Vorburger 2021). The idea is that with a greater variety of natural enemies each aphid hosts a different symbiont combination but is not protected against all attackers. Co-hosting symbionts can alter associated fitness costs (Zytynska, Tighiouart & Frago 2021), which can reduce aphid population growth rates or transmission of symbionts to the next generation (Rock *et al*. 2018). Additionally, the presence of multiple different symbiont combinations in the aphid population could alter competition among natural enemies (McLean & Godfray 2017), and as these selection pressures alter symbiont infection frequencies over time (Carpenter *et al*. 2021; Smith *et al*. 2021) this can further affect outcomes of hosting different symbiont combinations. Some symbionts may also be present due to non-natural enemy factors (host-plant or temperature mediated selection), leading to a strong context-dependency effect on interaction outcomes (Lemoine, Engl & Kaltenpoth 2020). Here, we aim to determine if different agricultural management practices can alter these selective pressures with potential consequences for natural pest control in crop systems.

Herbivore regulation by natural enemies is one natural ecosystem function that is often disrupted in managed landscapes, such as agroecosystems (Matson *et al*. 1997). The extent to which this is also exacerbated by insect protective symbionts is unknown. Many studies have shown the benefits of increasing plant diversity in agroecosystems (via banker plants or wildflower strips; Gurr *et al*. 2016; Tschumi *et al*. 2016; Balzan 2017), for pollinator and natural enemy populations, but often the effect of plant identity on specific plant-insect interactions is overlooked. Flowering plants offer variable sources of nectar, which is useful for adult insects whose offspring are the control agents (e.g. parasitoid wasps, lacewings, syrphids); parasitoid wasps can survive two weeks longer when a nectar source is offered, increasing search and attack rates (Russell 2015). These additional plants can also host populations of non-pest aphids on which to establish a diverse natural enemy population before pest aphids arrive on the crop plants. In a recent paper, we showed that aphid suppression was a result of numerous weaker interactions between different flower, pest, and natural enemy species, rather than a few dominant interactions (Zytynska *et al*. 2021), and here we used the same outdoor pot experiment to analyse the aphid symbiont communities alongside an additional large-scale field experiment.

We analysed bacterial symbiont communities hosted by cereal aphids (*Sitobion avenae* L.) and associations among these symbionts within aphids in two barley crop experiments: The first was a large-scale agricultural field experiment (Scotland), which compares long-term impacts of an integrated cropping system relative to standard commercial practice (Hawes *et al*. 2019) and the second an outdoor pot experiment (South Germany) where previous effects of flower identity on aphid control were demonstrated (Zytynska *et al*. 2021). Both experiments manipulated the presence of flowering plants, defined as the flower strip combined with the integrated approach for the field experiment or by the presence of three different flowering plant species for the pot experiment, across multiple barley varieties. The field experiment additionally examined the effect on aphid symbionts along a transect into the field, expecting decreased insect abundances into the fields (Thies & Tscharntke 1999) while the pot experiment examined the effects across time (Smith *et al*. 2015).

Our main hypotheses are (a) the presence of floral resources increases natural enemy abundance/diversity, which reduces aphid abundance, and (b) changes in natural enemy abundances and diversity alters the number, diversity and combination of symbionts hosted by aphids through changing selection pressures. We expect differences to be attributed to specific symbiont combinations as individual symbionts are not independent from one-another inside an aphid host (Mathé-Hubert *et al*. 2019); therefore, we examined effects across common symbiont combinations shared across multiple aphids. For the field experiment, only the total number of parasitoid wasps (via counts of aphid mummies, which host developing wasps) or general predators were collected (via insect traps) and thus we test the hypothesis only for parasitoid abundance rather than diversity. For the pot experiment, more comprehensive data was collected allowing for analysis of each type of parasitoid wasp and predators on aphids and their symbionts.

## Methods

### Large-scale agricultural field experiment

#### Study system and experimental design

The CSC platform at Balruddery Farm near Dundee, Scotland (described in Hawes *et al*. (2019)) is a 42-ha contiguous block of six arable fields, based on a six-year rotation of the commonly grown crops in the region. At the start of the first rotation in 2010, each of the six fields were divided in half. Conventional and integrated management treatments were randomly allocated to each half, and (for the barley fields) in each field-half three different barley varieties were planted as plot strips (i.e. 3 barley variety strips for conventional and 3 for integrated). The conventional treatment is the standard commercial management practice typical for the crop in terms of soil cultivation, fertiliser inputs and herbicide applications, while the integrated system aims to maintain yields, enhance biodiversity and soil biophysical quality, reduce non-renewable inputs and minimise losses from the system relative to conventional practice (Hawes *et al*. 2019). The integrated system includes organic amendment, conservation tillage, soil nutrient supply calculations to minimise use of mineral fertiliser, and reduced reliance on crop protection chemicals (IPM strategies, targeted weed control and species rich wildflower margins), Field margins around the integrated treatments were sown with a wildflower mix in 2015 (Balruddery Species Rich Meadow Margin, Scotia Seeds, UK containing seeds of 25 flower species).

*Sitobion avenae* aphids were collected on 6^th^ July 2016 and 3^rd^ July 2017, at four distances (5, 15, 30, 50 m) from the edge of the field into each barley variety strip (2016: *Cassata, Retriever, Saffron* and 2017: *Bazooka, Infinity, Retriever*). Aphids were stored in 70% ethanol and shipped to Technical University of Munich for symbiont analysis. At each sampling location (distance into the field strip), a maximum five aphids (one per colony) were collected from three adjacent infested tillers in every barley strip (variety); aphids reproduce asexually, and females deposit a group of offspring in one patch before moving away, this approach minimises the chance of collecting aphids from the same clonal mother. Total aphid number was also counted for these three tillers. In total, there were four sampling distances for three barley varieties, repeated across the two management systems, and across two years. Additional data were collected on % cover of the crop and weed plants in the sampled areas using a 1 m^2^ quadrat to estimate the projected area of ground covered by each vegetation type in mid-July. Data on parasitoid and generalist natural enemy abundance were obtained from two yellow sticky traps, one placed at the top and one at the bottom of the plant canopy, at each sampling location, in mid-to-late July.

#### Aphid endosymbiont analysis

DNA was extracted from individual aphids using the salting out method (Sunnucks & Hales 1996) and examined for nine common bacterial symbionts (*Hamiltonella defensa, Regiella insecticola, Serratia symbiotica, Rickettsia sp, Spiroplasma, Fukatsuia symbiotica, Rickettsiella viridis, Wolbachia*, and *Arsenophonus)* plus the primary symbiont (*Buchnera aphidicola*) as a positive control using PCR-based assays (Table S1) via gel electrophoresis (Zytynska *et al*. 2016). All samples were run alongside negative controls, with a subset repeated to ensure accuracy.

#### Data analysis

Data were analysed in R (v. 3.6.3) using RStudio (v. 1.2.5033). We analysed the data at two levels: the individual aphid (i.e. how many and which symbionts are hosted by individual aphids) and the local population level (i.e. how many and which symbionts are hosted across all aphids at each sampling location) where up to five aphids were sampled from three tillers per location.

We used generalised linear models to analyse the effect of year (2016, 2017), management system (conventional or integrated), distance in the field (5, 15, 30, 50 m), barley variety (Bazooka, Cassata, Infinity, Retriever, Saffron), weed cover and crop cover on overall aphid, parasitoid, and generalist predator abundance (quasipoisson error distribution for count data). Next, we used chi-square analyses to determine significant associations between symbiont species. A linear mixed effects model with year and barley variety as random effects was used to analyse the effect of management system, distance into field, crop cover, aphid abundance and interactions on the number of symbionts hosted by aphids (individual aphid and local population levels) and Shannon diversity of the aphid symbionts (local population level). Models were simplified by removing non-significant terms and minimum adequate models are presented. Further linear models were used to determine influences of management system and distance into the field on crop and weed cover (lmer, using asin transformed data for % cover producing a F-value statistic), and on parasitoid and predator abundances (glmer, using poisson error distribution for count data producing a Chi-square statistic). We analysed the effect of the management system, distance into the field and the abundance of parasitoid wasps (and interactions) on the proportion of aphids hosting different symbiont species (averaged across combinations) or the different symbiont combinations using generalised mixed effects models with binomial error distribution including year as a random effect. Lastly, we used Structural Equation Modelling (SEM) in the R package ‘piecewise’ (Lefcheck 2016), where responses included the proportion of aphids hosting each different symbiont combination, as well as aphid and parasitoid abundances, and relevant crop traits; predictors included experimental variables of management system and distance into the field. Model fit was evaluated using Fisher’s C statistic with the presented model reproducing the data well (P>0.05).

### Outside pot experiment

#### Study system and experimental design

The experiment was set up in Freising, Bavaria, South Germany on a paved surface next to a natural meadow, May-July 2018. We used four spring barley varieties that varied in aphid susceptibility in laboratory trials (Barke, Chevallier, Grace, and Scarlett) (Zytynska *et al*. 2020; Zytynska *et al*. 2021), and three flowering plant species commonly used as companion/intercropping plants with barley [Buckwheat (*Fagopyrum esculentum*), and Red clover (*Trifolium pratense*) from Rühlemann’s Kräuter & Duftpflanzen, Germany; and, Faba bean (*Vicia faba cv. Perla)*, from Kiepenkerl seeds].

We used a factorial randomised block experimental design that was focused on the effect of flower presence/absence and flower identity on aphid and natural enemy population dynamics on the barley (see Zytynska *et al*. 2021). The plants were placed in a grid system (4 × 5 plants per block), with 50 cm between each barley-flower combination, and 1.5 m between blocks. Here, we focus on *Sitobion avenae* aphids collected from barley plants that were grown either alone, or next to Fagopyrum, Trifolium or Vicia. The experiment also had a treatment with all three flower species (mixed) but aphid control was so effective here (Zytynska *et al*. 2021) that insufficient aphids could be collected for symbiont analysis. Seeds were germinated and grown in individual pots (2 litre pots filled with Floragard B Pot Medium-Coarse potting substrate, pH 5.6, NPK 1-0.6-1.2); one plant per pot for barley, Fagopyrum and Vicia was used while Trifolium plants were established from 15-20 sown seeds.

All plants were initially grown under an outdoor covered shelter for three weeks and then transferred fully outdoors on 29^th^ May 2018 and placed into individual 1 litre capacity pot trays. For the no flower treatment there was a single barley pot, for the treatments with one flower there was one barley pot and one flower pot. To avoid confounding factors related to ‘barrier effects’ of the plants, the flowers were placed behind the barley plants, away from the natural meadow (i.e. not between the barley and the meadow). Once per week, a full invertebrate survey was conducted by carefully examining every plant (both barley and flower) and recording the number of all generalist predators, specialist parasitoid wasp adults (if observed), and parasitoid mummies (hardened shell after an aphid has been parasitized, using form and colour to differentiate between mummies formed by *Aphidius sp, Aphelinus sp*. or *Praon sp*.). Aphids were identified to species, and winged/unwinged aphids were counted separately. Aphid and parasitoid DNA was amplified from a fragment of the insect CO1 gene using universal primers LCO1490 and HCO2198, to confirm species identification (Table S1). The dominant parasitoid species were identified as *Praon volucre* (pink mummies, with cocoon under the body) and *Aphidius rhopalosiphi* (copper-coloured mummies). No *Aphelinus* sp. parasitoid DNA was detected in any collected aphid samples, but black mummies (representative of this genus) were observed during the experiment.

#### Aphid endosymbiont analysis

Aphids were collected for endosymbiont analysis on 23^rd^ June (late June), 11^th^ July (early July) and 25^th^ July (late July). One aphid per colony on every plant was collected and stored in 70% ethanol. Only *Sitobion avenae* aphids were present in sufficient numbers for symbiont analysis. DNA extraction from whole aphids, and subsequent endosymbiont analyses using PCR-based assays (Table S1) were performed as for the large-scale experiment.

#### Data analysis

Data were analysed in R (v. 3.4.3) using RStudio (v. 1.0.143). We first analysed the effect of the presence and identity of the flowers, date of collection, as well as barley variety on overall *Sitobion avenae* aphid, parasitoid and generalist predator abundance using generalised linear models with quasipoisson error distributions. We also calculated natural enemy diversity using Shannon diversity and analysed the same factors as for abundance data but using a linear model with normal error distribution. Similarly, for the field experiment, we used Chi-square analyses to identify non-random associations among the symbiont species, important to interpret effects of symbiont combination as opposed to single individual symbiont species. Linear mixed effects models with experimental block and row as random factors were used to analyse the effect of date of collection, aphid abundance, barley variety, and flower treatment (and interactions) on the number of symbionts hosted by aphids (symbiont richness at the individual aphid and plant level, i.e. collected from the same plant) and on the Shannon diversity of symbionts at the plant level. We analysed the effect of symbiont species/combinations, flower treatment, and the natural enemy Shannon diversity and abundances of the different species (and interactions) on the proportion of aphids hosting different symbiont species or combinations (hosting or not hosting the symbiont) using generalised linear models with binomial error distribution. We also used Structural Equation Modelling (SEM) in the R package ‘piecewise’ (Lefcheck 2016), using linear mixed effect models with block and row as random effects. Responses included the proportion of aphids hosting each different symbiont combination, as well as abundance of aphids (winged and unwinged separated), predator (all species grouped together), and parasitoid (separated by genus); the number of non-pest aphids on the flowers was initially included but removed as it was non-significant in all models. Each aphid-symbiont combination was analysed using the number of aphids and natural enemies observed at the time point of collection, to account for differences in time of collection. Predictors included each flower resource as an individual variable to determine flower identity effects. Model fit was evaluated using Fisher’s C statistic with the presented model reproducing the data well (P>0.05).

## Results

### Effects of integrated management practices on aphid endosymbionts (CSC field experiment)

*Sitobion avenae* aphids (N=219) were collected from 41 sampling areas across the fields (termed local populations, 1-10 aphids per population, mean = 5.34 ± 0.29 SE, median/mode = 6). This included 144 aphids (71 from conventional, 73 from integrated) from 24 local populations in 2016, and 75 aphids (11 from conventional, 64 from integrated) from 17 local populations in 2017 from field strips that were managed according to either a standard conventional cropping practice in one field half or an integrated management system in the other field half (Hawes *et al*. 2019). We observed no significant effect of field management on total aphid abundance (F_1,36_=1.46, P=0.235; Fig. 1a) despite collecting fewer aphids from conventional fields in 2017 (but not 2016) – suggesting there were fewer but larger colonies in this year compared to integrated fields with more but smaller colonies (allowing for a greater number of aphids to be collected). We also observed no effect of management system on parasitoid wasp abundance (F_1,36_=0.71, P=0.403; Fig. 1b), but insect abundances decreased into the field-halves (aphids: F_1,36_=6.17, P=0.018, Fig. 1a; parasitoids: F_1,36_=8.08, P=0.007; Fig. 1b).

**Figure 1.**
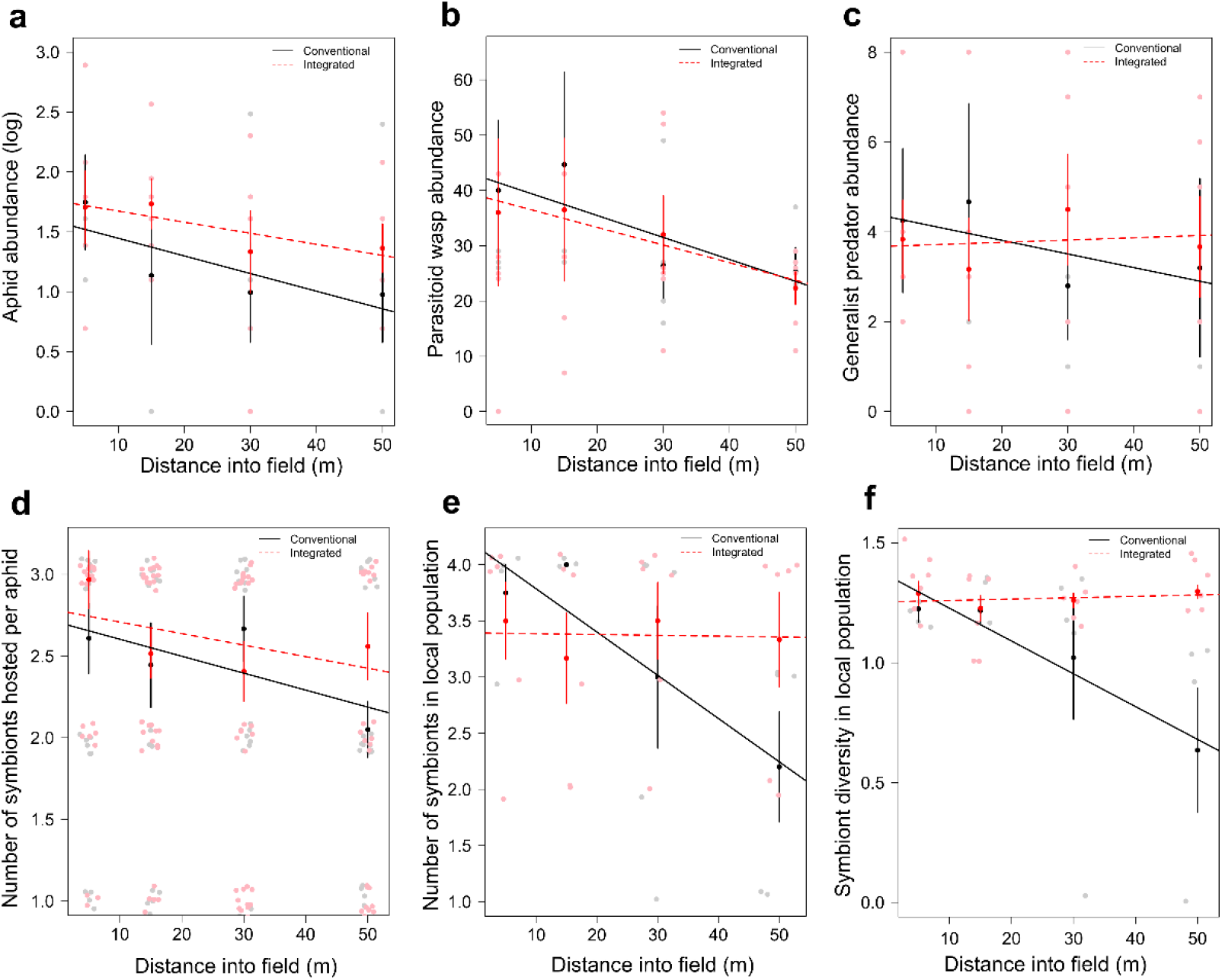
Effects of field management and distance into field on (a) aphid abundance (log transformed), (b) parasitoid abundance, (c) generalist predator abundance, (d) the number of symbionts hosted by individual aphids, (e) the number of symbionts within the local population (sampling area), and (f) the Shannon diversity of symbionts within the local population. Error bars represent ±1SE.

We observed no effect of management system or distance in the field for generalist predators (system: F_1,36_=0.12, P=0.730; distance: F_1,36_=0.02, P=0.895; Fig. 1c). Hence, our overall hypothesis that floral resources increase natural enemies and reduce aphids was not supported in this field system. However, across all field locations when there were more parasitoid wasps we observed fewer aphids (X^2^_1_=10.59, P=0.001), but the opposite with generalist predators where more predators were found in areas with higher aphid numbers (X^2^_1_=6.34, df=1, P=0.012). Lastly, integrated field-halves had reduced crop cover (F_1,36_=4.95, P=0.032) with correlated increases in weed cover (r= −0.65, n=41, P<0.001). Generalist predator abundance was positively associated with higher crop (X^2^_1_=13.81, P<0.001) and weed cover (X^2^_1_=3.84, P=0.05), with no significant effects on the abundance of specialist parasitoid wasps.

We identified four bacterial endosymbionts with high prevalence within the sampled aphids: *Hamiltonella defensa* (Hd) hosted by 63.9% of aphids; *Regiella insecticola* (Ri) hosted by 67.0% of aphids; *Serratia symbiotica* (Ss) hosted by 59.8% of aphids; and *Fukatsuia symbiotica* (Fs) hosted by 63.5% of aphids. Fifteen different combinations of these symbionts were identified in the aphids, with 20.1% of aphids (44/219) hosting all four. The most frequent symbiont combination included the three symbionts without *Serratia* (HdRiFs: 55/219, 25.1% of aphids), followed by aphids hosting all four symbionts and then those aphids singly hosting *Serratia* (Ss: 34/219, 15.5% of aphids). The average number of symbionts per aphid was 2.5 ± 0.1, while the average number of symbionts per sampling location (local population) was 3.3 ± 0.2. Fewer aphids (than expected at random) co-hosted any symbiont with *Serratia*, while more aphids co-hosted combinations of *Hamiltonella-Regiella* (X^2^_1_=36.0, df=1, P<0.001), *Hamiltonella-Fukatsuia* (X^2^_1_=34.6, df=1, P<0.001) and *Regiella-Fukatsuia* (X^2^_1_=50.1, df=1, P<0.001).

Aphids collected in integrated managed field strips co-hosted a slightly, but significantly higher number of symbionts (2.60 ± 0.09) than those in the conventional field-halves (2.45 ± 0.11), and symbiont richness per aphid decreased with distance into both field types by 18.6 % (Distance into field: Table 1, Fig. 1d). At the local population level, symbiont richness and diversity among aphids were also higher in the integrated fields (3.38 ± 0.18) than conventional (3.12 ± 0.28; Table 1). Symbiont species richness and diversity decreased into the field but only under conventional management, while remaining relatively constant into the field under integrated management (management system x distance into field, Table 1, Fig. 1e,f). The number of symbionts hosted by individual aphids strongly increased with increasing crop cover (Table 1), despite no effects of crop cover on aphid abundance (F_1,28_=0.01, P=0.927). With a crop cover of 20-60%, individual aphids hosted 1.85 ± 0.20 symbionts compared to 3.13 ± 0.12 symbionts with high crop cover of 65-95%. There was no effect of crop cover on the richness and diversity of symbionts at the local population level (Table 1).

**Table 1.** Effects of field management and distance into field on aphid symbiont communties.

|  | Symbiont richness |  |  |  |  |  | Symbiont diversity |  |  |
| --- | --- | --- | --- | --- | --- | --- | --- | --- | --- |
|  | Individual aphid level |  |  | Local population level |  |  | Local population level |  |  |
|  | df | F | P | df | F | P | df | F | P |
| Crop cover | 1,212 | **14.22** | **<0.001** | 1,34 | 0.20 | 0.730 | 1,34 | 3.69 | 0.066 |
| Aphid abundance | 1,212 | 0.16 | 0.686 | 1,34 | 1.63 | 0.210 | 1,34 | 0.29 | 0.593 |
| Parasitoid abundance | 1,212 | 0.29 | 0.590 | 1,34 | 0.04 | 0.852 | 1,34 | 1.19 | 0.282 |
| Management system | 1,212 | **8.94** | **0.003** | 1,34 | 0.74 | 0.396 | 1,34 | 0.01 | 0.986 |
| Distance into field | 1,212 | **9.33** | **0.003** | 1,34 | **5.19** | **0.029** | 1,34 | **7.83** | **0.009** |
| System x Distance | 1,212 | 2.42 | 0.121 | 1,34 | **7.59** | **0.009** | 1,34 | **9.70** | **0.004** |

*Linear mixed effects models (lmer) with barley variety and year as random effects. Distance into the field is* *fitted as a continuous variable.*

For further in-depth analysis of the endosymbiont combinations in aphids, we focus on the ten combinations that are hosted by a minimum of 5% of aphids within one management system (N=201 out of 219 aphids). The proportion of aphids hosting the different symbiont species varied across the management type (Χ^2^_1_=43.03, P<0.001, Fig. S1a) but not with the distance into the field (Χ^2^_3_=2.19, P=0.533, Fig. S1a). Aphids collected from conventional field-halves were more likely to host *Regiella*, while those from integrated field-halves were more likely to host *Serratia*. This is reflected in the common combinations of symbionts hosted by aphids (combination x management system: Χ^2^_9_=26.23, P=0.002; combination x distance: Χ^2^_9_=8.41, P=0.493) (Fig. 2a). Here, it can also be seen that aphids collected from conventional field-halves host (on average) combinations with fewer symbionts than aphids from integrated ones, with more aphids hosting 1-2 symbionts in the inner part of conventional fields (Fig. 2a).

**Figure 2.**
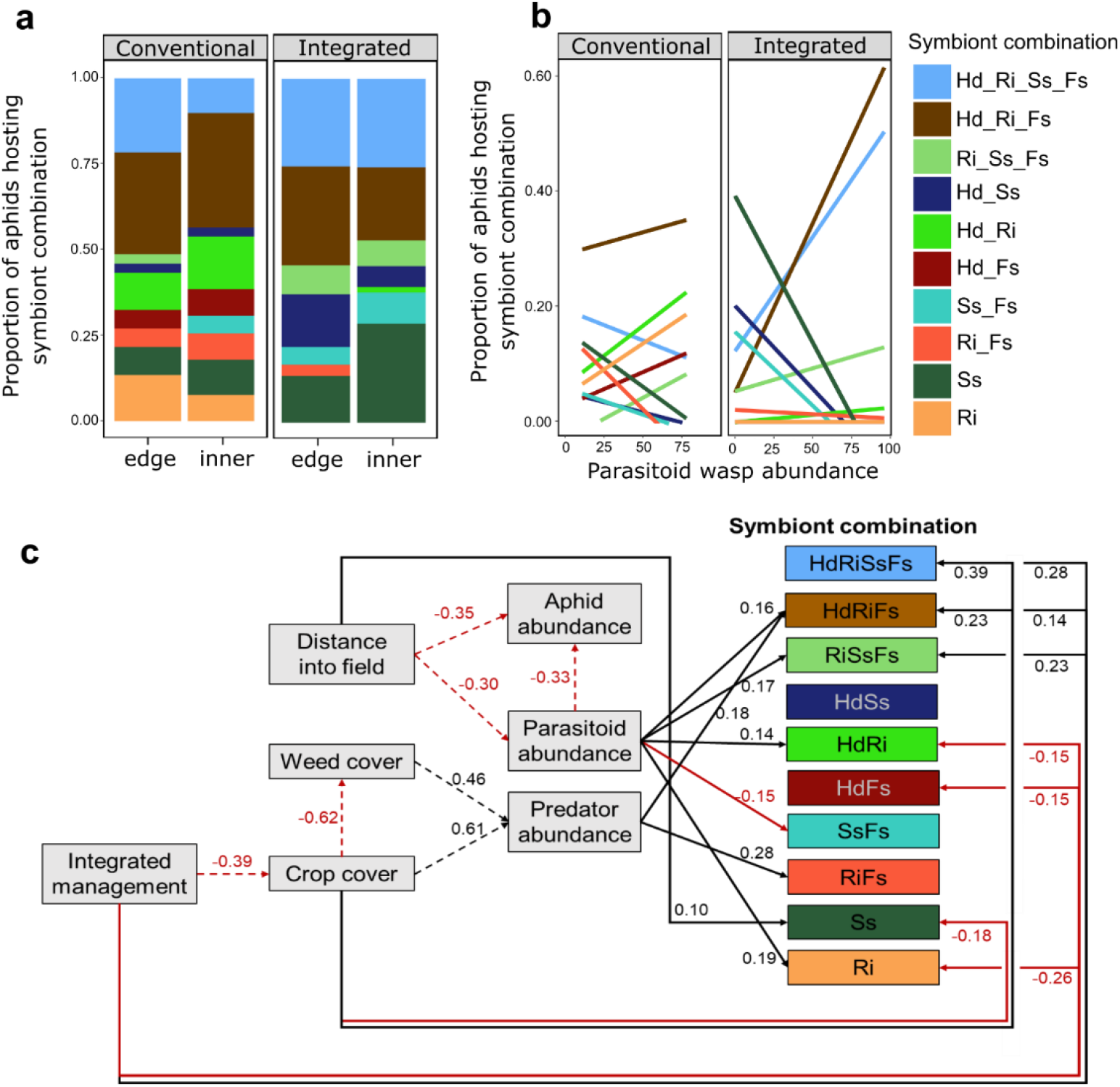
Field experiment: effects of field variables on the aphid symbiont community. The proportion of aphids hosting the ten most common most common combinations of symbionts within integrated and conventionally managed fields across (a) distance into the field (edge, 5-15m or inner, 30-50m), and (b) parasitoid wasp abundance, highlighting variation due to symbiont combinations as opposed to overall symbiont richness or diversity (Table 1). Further explored in (c) via structural equation modelling showing significant links between variables and symbiont combinations. Dashed lines show analysis at the level of sampling location, with solid lines showing results at the individual aphid level. *Serratia symbiotica* (Ss), *Rickettsia* (Rk), *Wolbachia* (W), *Regiella insecticola* (Ri), *Hamiltonella defensa* (Hd) and *Spiroplasma* (Sp).

The abundance of parasitoid wasps in the local sampling area influenced the number of aphids hosting the different symbiont species (Χ^2^_3_=50.87, P<0.001; Fig. S1b) or combinations (Χ^2^_9_=48.84, P<0.001) (Fig, 2b). This was also dependent on the management system (Χ^2^_2_=20.77, P<0.001; Fig. 2b) but not the distance into the field, perhaps due to the mobility of the parasitoids. Using structural equation modelling (Fig 2c), we show that there are direct positive and negative effects of integrated management as well as parasitoid and predator abundances on the number of aphids hosting specific symbiont combinations. Indirect effects via the distance into the field, and effects of management on crop and weed cover further mediate effects of natural enemies on the number of aphids hosting the different symbiont combinations. The level of protection provided by these different symbiont combinations against natural enemies remains to be tested.

### Flower identity effects on aphid symbionts (outdoor pot experiment) Outside pot experiment

Aphid abundance was reduced on barley plants by 30 % when grown next to a flower (F_1,118_=10.06, P=0.002), with fewest aphids on barley grown next to Fagopyrum or Trifolium (F_2,93_=3.92, P=0.024; Fig 3a). Parasitoid wasp abundance varied strongly with flower treatment and identity (F_3,116_=5.76, P=0.001) and was lowest on barley next to no flower while highest on barley next to Trifolium (120% higher, Fig. 3b), with effects on overall natural enemy diversity incorporating parasitoid wasp species as well as generalist predators such as ladybird and lacewing larvae primarily driven by the presence of flowers more than the identity (F_1,94_=3.78, P=0.055, Fig. 3c). The abundance of *S. avenae* aphids was higher on barley varieties Grace and Scarlett than on Barke or Chevallier (F_3,110_=3.33, P=0.022), while there were no differences across barley varieties in parasitoid abundances (F_3,91_=0.85, P=0.467) or natural enemy diversity (F_3,91_=1.43, P=0.239).

**Figure 3.**
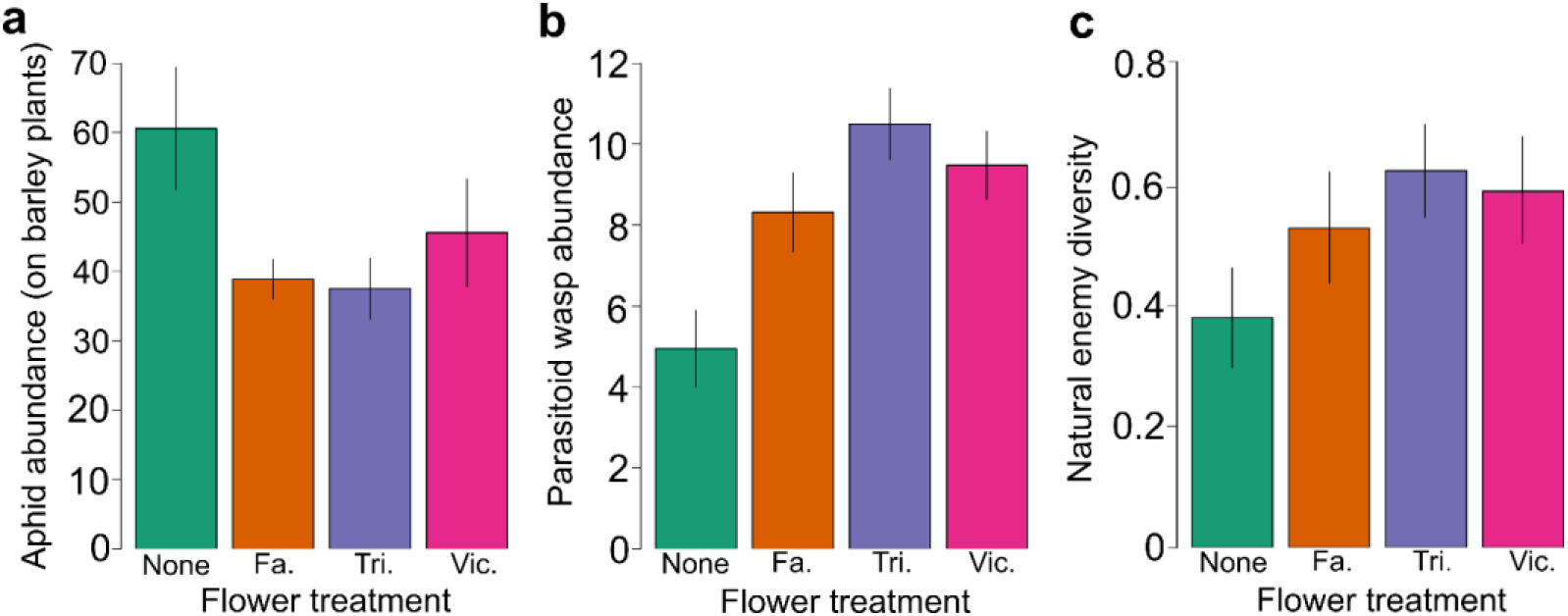
Outdoor pot experiment: effects of flower treatment on (a) overall aphid abundance, (b) parasitoid wasp abundance, (c) natural enemy diversity. None: no flower, Fa: *Fagopyrum esculentum*, Tri: *Trifolium pratense*, Vic: *Vicia faba*. Error bars represent ± 1SE.

For symbiont analysis,127 *Sitobion avenae* aphids were collected from barley plants grown alone (N=34), next to Fagopyrum (N=33), Trifolium (N=23) or Vicia (N=37) plants, with 29-36 aphids collected from each barley variety across flower treatments. Of these aphids, we collected 63 in late June, 43 in early July and 21 in late July. We identified six bacterial symbionts within the sampled aphids: *Serratia symbiotica* (Ss) was hosted by all sampled aphids (100%), *Rickettsia* (Rk) and *Wolbachia* (W) each were hosted by 77/127 (60.6%) aphids, *Regiella insecticola* (Ri) and *Hamiltonella defensa* (Hd) were hosted by 72/127 (56.7%) aphids while *Spiroplasma* (Sp) was hosted by 54/127 (42.5%) aphids. Thus, all aphids hosted at least one symbiont and 8 (6.3 %) aphids hosted all 6 symbionts. There were 29 different combinations of symbionts observed across all aphids, however only eight combinations were each hosted by at least 5% of the population and these are the focus for the analyses (N=84 aphids of 127 total aphids with symbionts). The remaining 21 less abundant combinations had an average infection frequency of 2.0 ± 0.29 %. The most frequent symbiont combinations were the 4-symbiont combination of SsRkHdW (10.2%) and RiSsRkW (8.7%). There was a positive association between frequencies of symbionts *Wolbachia* and *Regiella* (Χ^2^_1_=5.41, P=0.020), and between *Wolbachia* and *Rickettsia* (Χ^2^_1_=24.50, P<0.001), while we detected a negative association between *Regiella* and *Hamiltonella* (Χ^2^_1_=21.46, P<0.001).

The number of symbionts co-hosted by individual aphids, and the number and diversity across all aphids collected from the same plant (local population level) was strongly influenced by the date of collection and the overall abundance of aphids on each plant (Table 2). The number of symbionts hosted by aphids increased from late June (3.56±0.14) to early July (4.26±0.17), and then dropped again towards the end of July (3.43±0.17). Flower identity influenced the number of symbionts hosted by individual aphids and at the population level (Table 2), with aphids hosting fewer symbionts when collected from barley grown next to Trifolium or Vicia. At the population level, barley variety was not a significant main effect for symbiont species richness or diversity. Nevertheless, the effect of the flower treatment was dependent on the barley variety driven by a strong decrease in the number of symbionts hosted by aphids on *Grace* barley next to Trifolium.

**Table 2.** Effects of flower presence and identity on aphid symbiont communties.

|  | Symbiont richness |  |  |  |  |  | Symbiont diversity |  |  |
| --- | --- | --- | --- | --- | --- | --- | --- | --- | --- |
|  | Individual aphid level |  |  | Plant level |  |  | Plant level |  |  |
|  | df | F | P | df | F | P | df | F | P |
| Date of collection | 2,107 | **7.27** | **<0.001** | 2,56 | **8.91** | **<0.001** | 2,56 | **6.79** | **0.002** |
| Aphid abundance | 1,107 | **4.04** | **0.047** | 1,56 | **5.12** | **0.028** | 1,56 | **4.79** | **0.038** |
| Nat. enemy diversity | 1,107 | 0.13 | 0.717 | 1,56 | 2.57 | 0.114 | 1,56 | **4.24** | **0.044** |
| Barley variety | 3,107 | 1.30 | 0.278 | 3,56 | 1.83 | 0.152 | 3,65 | 2.44 | 0.074 |
| Flower PA | 1,107 | 0.72 | 0.397 | 1,56 | 0.11 | 0.741 | 1,56 | 0.03 | 0.873 |
| Flower identity | 3,107 | **3.35** | **0.039** | 3,56 | **3.71** | **0.031** | 1,56 | 2.68 | 0.078 |
| Barley x Flower id | 9,107 | 0.87 | 0.551 | 9,56 | **2.25** | **0.032** | 3,56 | **2.25** | **0.031** |

*Linear mixed effects models (block and row as random), aphid number is by date of collection. No significant interactions with date of collection were observed.*

Flower identity strongly affected the symbiont combinations hosted by cereal aphids on the adjacent barley plants. The proportion of aphids hosting each symbiont species (Χ^2^_5_=25.37, P=0.045; Fig. S2) and combination of symbionts (Χ^2^_7_=42.43, P=0.004; Fig. 4a) differed among the flower treatments. Aphids collected next to Trifoilum and Vicia (the legumes) did not host the 6-symbiont combination of symbionts, and no aphids hosting the 2-symbiont combination of *Regiella-Serratia* were collected in the Fagopyrum treatment. The date of collection altered the frequency of aphids hosting the different symbiont combinations (Χ^2^_2_=65.30, P<0.001; Fig. 4b). Of note, the six-symbiont combination plus the 5-symbiont combination without *Hamiltonella* was hosted by half of all aphids collected in early July (peak aphid and natural enemy abundances), with a strong decrease in these abundances by late July.

**Figure 4.**
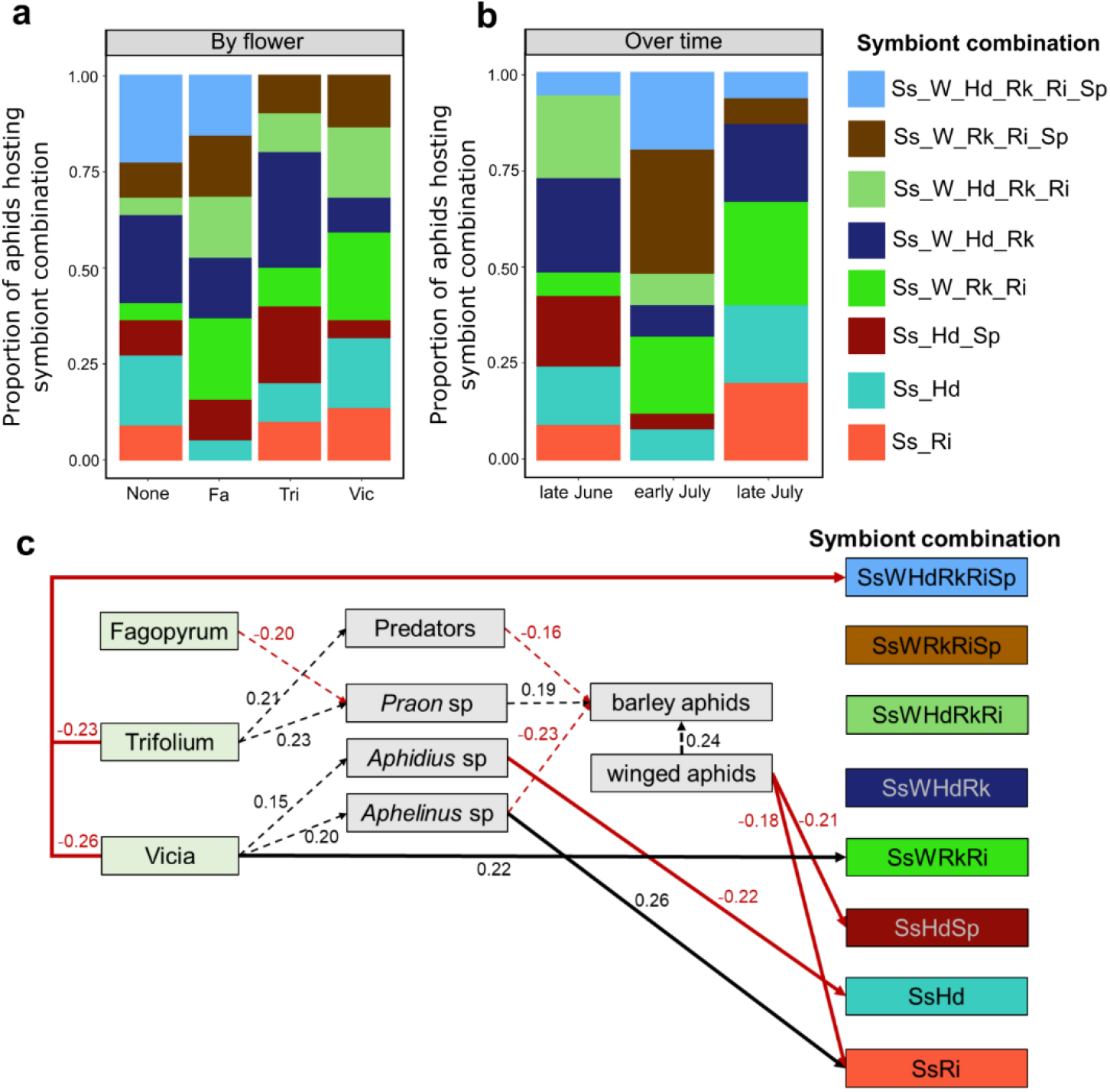
Outdoor pot experiment: the effect of flower treatment and time on aphid symbiont communities. The proportion of aphids hosting the eight most common symbiont combinations (a) across flower treatments, and (b) over time, highlighting variation on specific combinations of symbionts as well as on overall richness and diversity effects (Table 1). Further explored in (c) via structural equation modelling showing significant links between variables and symbiont combinations, using data combined across all dates but incorporating appropriate abundance data relating to aphid collection date (see methods). Dashed lines show analysis at the level of plant, with solid lines showing results at the individual aphid level. *Serratia symbiotica* (Ss), *Rickettsia* (Rk), *Wolbachia* (W), *Regiella insecticola* (Ri), *Hamiltonella defensa* (Hd) and *Spiroplasma* (Sp).

Lastly, we found that the number of aphids hosting the dominant symbiont combinations varied dependent on the flower treatment and the natural enemy community recruited by the different flowers (Fig. 4c). Using structural equation modelling, we were able to highlight direct and indirect effects of flower identity on the number of aphids hosting the various symbiont combinations. For example, Vicia plants led to an increase in *Aphidius sp*. parasitoid wasps that were negatively associated with aphids hosting *Serratia-Hamiltonella* symbionts, and an increase in *Aphidius sp*. parasitoids that were positively associated with aphids hosting *Serratia-Regiella* symbionts (Fig. 4c). We found that natural enemy diversity did not correlate with overall symbiont diversity, except in the Trifoilum treatments (Fig S3), but rather via abundances of specific individual symbiont combinations (Fig. 5). By analysing correlations between natural enemy diversity and the proportion of aphids hosting the different common symbiont combinations (Χ^2^_7_=36.13, P=0.021; Fig. 5a) we identified those symbiont combinations most associated with changes in natural enemy diversity and abundance of each natural enemy group (Fig. 5b-e). While the level of protection provided by these different symbiont combinations against natural enemies remains to be tested, we can highlight some interactions worthy of future focus: a consistent increase in aphids hosting symbiont combination *Serratia-Hamiltonella-Spiroplasma* (SsHdSp) was seen with increasing *Aphidius* sp. parasitoid wasp abundance while those hosting SsHd decreased (Fig. 5b); *Aphelinus* sp. parasitoids were not present on barley plants next to no flower, but for those next to Trifoilum or Vicia more aphids hosting *Regiella-Serratia* (RiSs) were collected when parasitoid abundance was higher (Fig. 5c).

**Figure 5.**
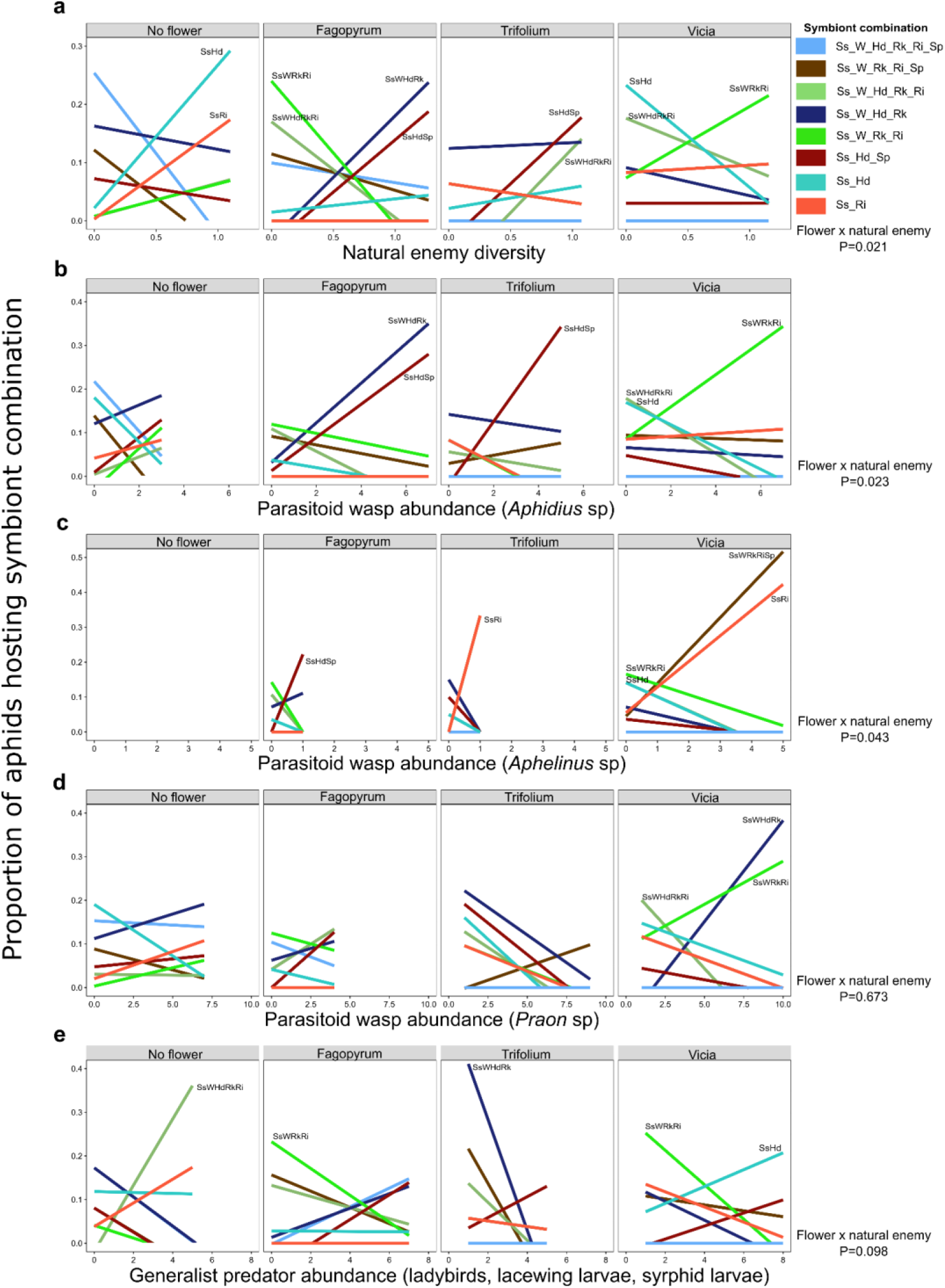
Outdoor pot experiment: summary of associations between natural enemy abundance and aphid symbiont communities across the flower treatments. The proportion of aphids hosting the eight most common symbiont combinations across (a) natural enemy Shannon diversity, and abundance of (b) *Aphidius* sp. (c) *Aphelinus* sp. (d) *Praon* sp. parasitoid wasps and (e) generalist predators. Statistical significance of interaction shown to the right. *Serratia symbiotica* (Ss), *Rickettsia* (Rk), *Wolbachia* (W), *Regiella insecticola* (Ri), *Hamiltonella defensa* (Hd) and *Spiroplasma* (Sp).

## Discussion

The outdoor pot experiment showed a clear reduction in aphids on barley plants in the presence of flowers, with correlated increases in natural enemy abundance and diversity; the mixed flower treatment suppressed aphids so strongly that insufficient aphids could be collected for endosymbiont analyses (further explored in Zytynska *et al*. (2021)). Flower identity affected the type of natural enemy recruited (species of parasitoid wasp or combined generalist predators) and abundances of these correlated with the proportion of aphids hosting different endosymbiont communities. Variation across time further demonstrated the importance of temporal changes in aphid and natural enemy abundances on endosymbiont communities, also shown in pea aphid systems by Smith *et al*. (2015) and to a limited extent by Leclair *et al*. (2021). The large-scale field experiment comparing aphids collected from field plot strips grown under an integrated or conventional management system (Hawes *et al*. 2019) did not result in reduced aphid or increased natural enemy abundances in integrated-managed fields. However, in accordance with the pot experiment we again identified a range of associations between natural enemy abundances and the proportion of aphids hosting specific endosymbiont combinations. There were further links identified between integrated management and crop cover on aphid endosymbiont communities independent of natural enemy variables, indicating additional effects beyond those studied here. Overall, we observed aphids co-hosting a greater number of endosymbionts in integrated field strips, while in the pot experiment this effect was dependent on the natural enemy abundance and driven by effects with Trifoilum plants. This highlights the context-dependency aphid-endosymbiont-natural enemy effects, but suggests management changes could have immediate effects on the surrounding ecological community with potential within-season impacts for pest control of cereal aphids.

In the field experiment, aphids collected in the integrated field-strips hosted higher numbers of endosymbionts (3- and 4-species combinations). While our study did not empirically test for symbiont-induced costs or the protective benefits of hosting these symbionts, we infer that correlations with parasitoid abundances indicate protective effects: aphids surviving in areas of higher parasitism are more likely to host protective symbionts. The costs of co-hosting multiple symbionts also remain to be tested for these specific aphids and symbionts, but since aphid abundances did not differ between conventional and integrated systems this may indirectly indicate increased hosting costs. Previous work, predominantly on other aphid species, has shown that parasitoid resistance can be conferred by all the symbionts we identified (reviewed by Guo *et al*. 2017). In *Sitobion avena*e aphids experimental work with endosymbionts has shown *R. insecticola* to reduce parasitoid emergence rates (Luo *et al*. 2020) while infection with *H. defensa* had no protective effect (Lukasik *et al*. 2013; Zepeda-Paulo, Villegas & Lavandero 2017; Li *et al*. 2018). Effects of the other endosymbionts has not been empirically tested in *S. avenae* sufficiently to make conclusions, but our results suggest that examining variation between *Serratia symbiotic* and *R. insecticola* would be an informed starting point. Future work should also focus on naturally occurring combinations of symbionts as opposed to single infections since these drive the responses we identified rather than individual effects of one symbiont species.

In the pot experiment, aphid abundance was reduced and natural enemy diversity/abundance increased on barley plants grown next to a flower through effects on non-pest aphids on the flowers before aphid colonisation on the adjacent barley (also see Zytynska *et al*. 2021). Thus, our experiment confirms effects of wildflower strips seen in small scale experiments (Tschumi *et al*. 2016; Balzan 2017; Albrecht *et al*. 2020), and indicates a need for more in-field (close-range) floral resources to maximise effects; for example, via intercropping or in-field floral rows. We observed significant variation in the symbionts hosted by aphids due to flower identity as well as presence, and these correlated with abundances of the different natural enemies recruited. These correlations can inform future studies to identify specific costs and benefits, which will allow a greater understanding of how this trade-off differs among symbiont combinations to alter aphid population dynamics. Of particular interest are the correlations of multi-symbiont combinations SsWRkRi, SsHdSp and SsWHdRk with *Aphidius* parasitoids and SsRi with *Aphelinus* parasitoids, again highlighting the potential importance of *R. insecticola* (Ri) in these aphids (Luo *et al*. 2020). In our previous study (Zytynska *et al*. 2021) we highlighted interactions among the different natural enemies within flower treatment and these could have impacts also for the aphid endosymbiont community. Parasitoid wasps avoid attacking aphids that they detect to have been previously attacked, with this effect lasting up to two days (Outreman *et al*. 2001). This will increase the survival chance of aphids resistant to the original parasitoid but perhaps not the second attacker, especially if it is a different species. In contrast, generalist predators readily consume parasitized aphids before mummification (Meisner *et al*. 2011), with some evidence that *R. insecticola* symbionts may also increase predation risk of aphids by ladybirds (Ramírez-Cáceres *et al*. 2019). Predation of parasitised aphids removes the parasitoid from the next generation via intraguild predation, thus the effect on biocontrol will be dependent on the local aphid, symbiont, and natural enemy community composition. The rare but ecologically-important potential of horizontal transmission of symbionts among aphids via natural enemies (Oliver *et al*. 2010; Gehrer & Vorburger 2012; Zytynska & Venturino 2018) may further explain the ability of aphid populations to response rapidly to changing natural enemy abundances. Overall, this leads to complex community dynamics and diverse selection pressures acting on the symbiont communities, particularly where there is high natural enemy diversity; any effect of aphid or parasitoid genotype (not studied here) can further influence outcomes. We also observed effects of the number of winged aphids on the proportion of unwinged aphids hosting 2- and 3-symbiont combinations. Winged aphids are long-distance dispersal morphs and could be important if they bring in novel symbiont combinations, assuming reproduction within the crop, but few studies have so far assessed symbionts in winged aphids (but see Smith *et al*. (2021)). Identifying source populations in the wider ecological landscape, e.g. from surrounding cereal fields, will help us understand community dynamics and potential effects on local biological control efforts.

Our hypothesis was that a low diversity of natural enemies favours selection for a combination of symbionts that provides the highest level of protection (Oliver *et al*. 2008) and subsequent extinction of the natural enemy (Sanders *et al*. 2016; Vorburger 2018). From our data, a dominance of *Aphelinus* parasitoids might select for aphids hosting *Serratia-Regiella*, while a dominance of *Aphidius* parasitoids might select for those hosting *Serratia-Hamiltonella-Spiroplasma*. In a diverse system (e.g. with floral resources increasing natural enemy diversity), there will be a mixture of these natural enemies leading to a mixture of symbiont combinations.

Variable effects of different natural enemies and interactions among them is key to disrupting selection for symbiont combinations with high protective effects but low fecundity costs (Zytynska & Meyer 2019). We showed this does not necessarily translate to increased diversity of aphid symbionts in a local population but rather that it is specific to the symbiont combinations that are hosted and the interactions experienced by the population. When every interaction differs for the various natural enemy species this can contribute to natural regulation of pest populations (e.g. Preedy *et al*. 2020), by naturally diversifying biological control strategies for crop protection (Pimentel 1991).

In conclusion, we show that floral plantings, and flower identity, can have community-wide effects on the diversity of natural enemies of herbivores, aphid populations, and the bacterial endosymbionts hosted by the aphids. Management of agricultural systems via conservation biological control and regenerative approaches requires ecological knowledge to predict how these might impact other parts of the system to optimise yield outputs. Unravelling what drives herbivore population regulation enables us to understand the important role of complex multi-species interactions and highlights the processes we should aim to promote to enhance natural pest regulation. Designing tailored floral plantings (e.g. Tschumi *et al*. 2016) to contain sufficient functional diversity to recruit and establish an abundant and diverse community of natural enemies (via nectar and non-pest prey) is key to increasing their impact for pest control on neighbouring crops. A greater diversity of natural enemies is likely to reduce in-field selection pressures favouring protective endosymbiont communities of pest insects, which might otherwise undermine natural pest regulation, and suggests a mechanism by which diversity can be utilised for internal system regulation, reducing reliance on agrochemical inputs for pest control.

## Author contributions

SEZ, AK, CH designed the study and collected the data. SS performed the molecular analyses to identify the symbionts. SEZ analysed the data, WWW supported the interpretation of results. SEZ wrote the manuscript with comments from all authors.

## Acknowledgements

We thank the CSC Balruddery team for management of the field experiment (https://csc.hutton.ac.uk). SEZ was supported by a British Ecological Society small research grant (SR16/1069) and a BBSRC (UKRI) David Phillips Fellowship BB/S010556/1. AJK and CH were supported by the Scottish Government’s strategic research programme funded by the Rural & Environment Science & Analytical Services Division.

