## Supplementary information for "Floral presence and flower identity alter cereal aphid endosymbiont communities on adjacent crops"

*[1] Department of Evolution, Ecology, and Behaviour. Institute of Infection, Veterinary and Ecological Sciences. University of Liverpool, Crown Street, Liverpool, L69 7ZB, UK*

*[2] Technical University of Munich, Terrestrial Ecology Research Group, Department of Life Science Systems, School of Life Sciences, Hans-Carl-von-Carlowitz-Platz 2, 85354 Freising, Germany*

*[3] Ecological Sciences Department, The James Hutton Institute, Invergowrie, Dundee, DD2 5DA, UK*

**Table S1: Primers, and sequences used to identify endosymbionts of cereal aphids**

**Fig S1. Field experiment: effects of field variables on the aphid symbiont species**

**Fig S2. Outdoor pot experiment: the effect of flower treatment on aphid symbiont species**

**Fig S3. Outdoor pot experiment: the relationship between natural enemy diversity and symbiont diversity, over flower treatment and time.**

**Table S1: Primers, and sequences used to identify endosymbionts of cereal aphids**

| Symbiont Species | Forward | sequence | Reverse | sequence |
|---|---|---|---|---|
| *Buchnera symbiotica* | Buch16S1F | GAGCTTGCTCTCTTTGTCGGCAA | Buch16S1R | CTTCTGCGGGTAACGTCACGAA |
| *Hamiltonella defensa* | 10F | AGTTTGATCATGGCTCAGATTG | T419R | AAATGGTATTCGCATTTATCG |
| *Regiella insecticola* | 10F | AGTTTGATCATGGCTCAGATTG | U443R | GGTAACGTCAATCGATAAGCA |
| *Serratia symbiotica* | 10F | AGTTTGATCATGGCTCAGATTG | R443R | CTTCTGCGAGTAACGTCAATG |
| *Fukatsuia symbiotica* | 10F | AGTTTGATCATGGCTCAGATTG | X420R | GCAACACTCTTTGCATTGCT |
| *Rickettsia* | 16SA1 | AGAGTTTGATCMTGGCTCAG | Rick16SR | CATCCATCAGCGATAAATCTTTC |
| *Spiroplasma* | 10F | AGTTTGATCATGGCTCAGATTG | TKSSsp | TAGCCGTGGCTTTCTGGTAA |
| *Rickettsiella* | RCL16S-211F | GGGCCTTGCGCTCTAGGT | RCL16S-470R | TGGGTACCGTCACAGTAATCGA |
| *Wolbachia* | W-SpecF | CATACCTATTCGAAGGGATAG | W-SpecR | AGCTTCGAGTGAAACCAATTC |
| *Arsenophonus* | Ars-23S1F | CGTTTGATGAATTCATAGTCAAA | Ars-23S2R | GGTCCTCCAGTTAGTGTTACCCAAC |
| aphid COI | LCO1490 | GGTCAACAAATCATAAAGATATTGG | HCO2198 | TAAACTTCAGGGTGACCAAAAAAT |

| **Master Mix** | µl per reaction |
|---|---|
| DNA | 2.0 |
| 5x Bioline MyTaq Reaction Buffer | 4.0 |
| PrimerF (20µM) | 0.4 |
| PrimerR (20µM) | 0.4 |
| Bioline MyTaq (5U/µl) | 0.2 |
| $ddH_2O$ | 13.0 |
| total | 20.0 |

| **Touchdown PCR program 65-55°C** | | |
|---|---|---|
| 94°C | 05:00 | |
| 94°C | 00:15 | 10x temp **-1°C/**cycle |
| 65-55°C | 00:30 | |
| 72°C | 00:30 | |
| 94°C | 00:15 | 25x |
| 55°C | 00:30 | |
| 72°C | 00:30 | |
| 72°C | 06:00 | |
| 4°C | ∞ | |

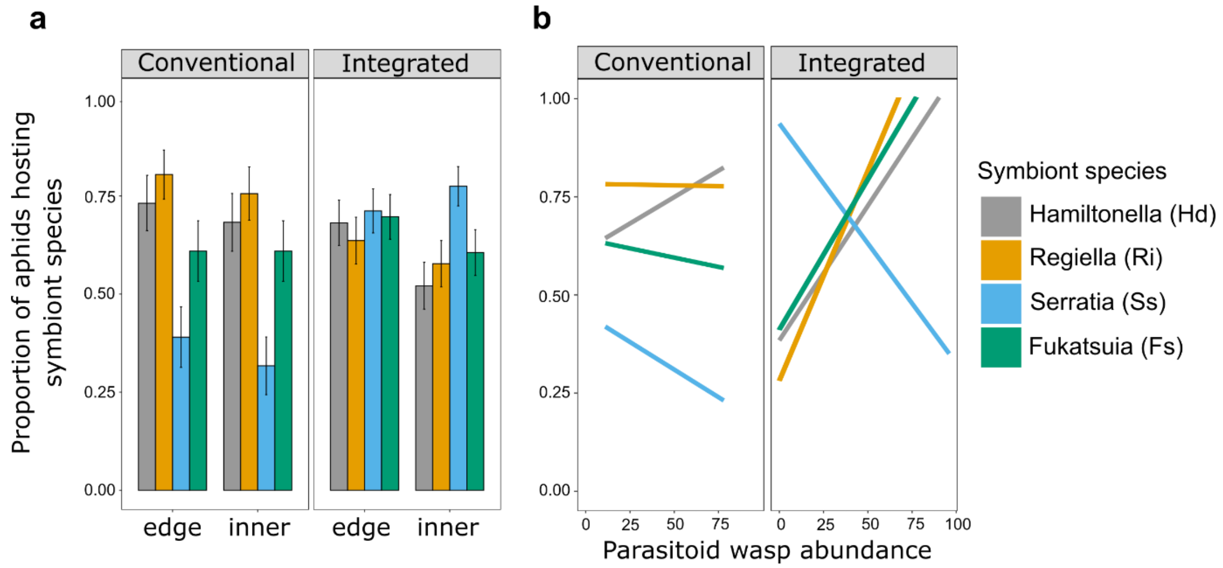


**Fig S1. Field experiment: effects of field variables on the aphid symbiont species.** The proportion of aphids hosting the different symbiont species within integrated and conventionally managed fields across (a) distance into the field (edge, 5-15m or inner, 30-50m), and (b) parasitoid wasp abundance. Aphid host multiple endosymbionts and therefore sum of proportions will be more than one. Error bars represent ±1SE.

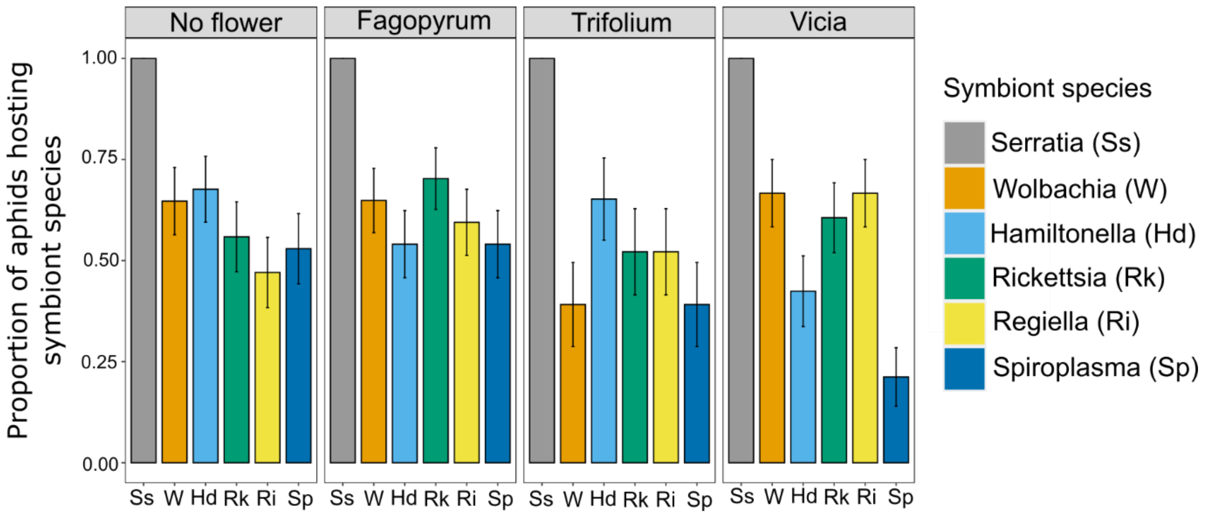


**Fig S2. Outdoor pot experiment: the effect of flower treatment on aphid symbiont species.** The proportion of aphids hosting different symbiont species across flower treatments. Error bars represent ±1SE.

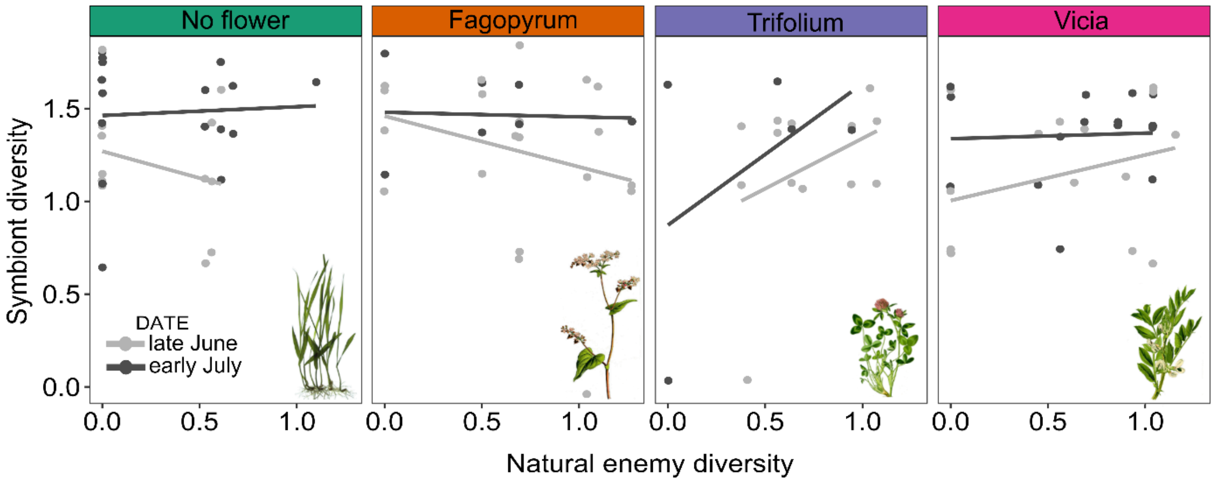


**Fig S3. Outdoor pot experiment: the relationship between natural enemy diversity and symbiont diversity, over flower treatment and time.** Only data for late June (n=71 aphids on 50 plants) and early July (n=53 aphids on 32 plants) shown in (c) due to lack of sufficient data for comparisons in late July (n=21 aphids on 12 plants).
